# An attention-based deep neural network model to detect cis-regulatory elements at the single-cell level from multi-omics data

**DOI:** 10.1101/2024.10.20.619317

**Authors:** Ken Murakami, Keita Iida, Mariko Okada

**Affiliations:** Laboratory for Cell Systems, Institute for Protein Research, Osaka University, Suita, Japan; Integrated Frontier Research for Medical Science Division, Institute for Open and Transdisciplinary Research Initiatives (OTRI), Osaka University, Suita, Japan

**Keywords:** Cis-regulatory elements, Enhancers, Gene regulation, Deep learning, Attention-based neural network, Explainable artificial intelligence, Single-cell multiome, Single-cell ATAC-seq, Single-cell analysis, Intra-tumor heterogeneity

## Abstract

Cis-regulatory elements (cREs) play a crucial role in regulating gene expression and determining cell differentiation and state transitions. To capture the heterogeneous transitions of cell states associated with these processes, detecting cRE activity at the single-cell level is essential. However, current analytical methods can only capture the average behavior of cREs in cell populations, thereby obscuring cell-specific variations. To address this limitation, we proposed an attention-based deep neural network framework that integrates DNA sequences, genomic distances, and single-cell multi-omics data to detect cREs and their activities in individual cells. Our model shows higher accuracy in identifying cREs within single-cell multi-omics data from healthy human peripheral blood mononuclear cells than other existing methods. Furthermore, it clusters cells more precisely based on predicted cRE activities, enabling a finer differentiation of cell states. When applied to publicly available single-cell data from patients with glioma, the model successfully identified tumor-specific SOX2 activity. Additionally, it revealed the heterogeneous activation of the ZEB1 transcription factor, a regulator of epithelial-to-mesenchymal transition-related genes, which conventional methods struggle to detect. Overall, our model is a powerful tool for detecting cRE regulation at the single-cell level, which may contribute to revealing drug resistance mechanims in cell sub- populations.

## Introduction

The human genome contains approximately 20,000 protein-coding genes and over 900,000 cis-regulatory elements (cREs) that regulate gene expression (Moore et al., 2020). Typical cREs, known as enhancers, contain transcription factor (TF) binding sites and are located at distances ranging from a few kilobase pairs to several megabase pairs from the transcription start site (TSS) (Panigrahi et al., 2021). Enhancers make physical contact with promoters via TFs and are known to activate gene expression, regardless of their orientation (Yang et al., 2024). Enhancers act as on/off switches for gene expression, and variations in active enhancer regions contribute to differences in cell types and species (Villar et al., 2015). Although the coding regions follow clear rules, as determined by the DNA codon table, the regulation of cREs is highly complex. Active cREs are typically present in an open chromatin state (Thurman et al., 2012) and regulate gene expression by allowing TFs to bind to specific motifs (Spitz et al., 2012). However, the cell controls when and where the genome becomes accessible, as well as the number of TFs that can bind cREs, depending on the cell state. In addition, stochastic changes in chromatin accessibility are known to exist within heterogeneous in cell populations (Bohrer et al., 2021). This heterogeneity is thought to be a driving force for cell state transition, as shown in the Waddington Landscape (Waddington, 1957), and contributes to the plasticity and drug resistance of cancer cells in pathological contexts (Marusyk et al., 2020). Therefore, to elucidate transcriptional regulation and the control of multicellular processes, detecting cREs and their effects on gene expression levels in individual cells is essential.

A single-cell assay for transposase-accessible chromatin sequencing (scATAC-seq) (Buenrostro et al., 2015) has been widely used to detect the open chromatin regions at the single-cell level. Several computational methods have been developed to annotate functions and detect cREs in scATAC-seq-derived open chromatin regions. Early approaches to detect cRE-gene relationships from single-cell data were based on the correlation between the chromatin accessibility of cRE candidates and the gene expression level (or chromatin accessibility at the promoter) of target genes (Pliner et al., 2018; Granja et al., 2021). Despite being significantly faster at detecting cREs compared to traditional reporter assays, these correlation-based methods also detect false-positive pairs, especially when the dataset does not contain a sufficient variety of cell states or cell types. To address this issue, curation rules such as penalization were introduced, based on the genomic distance from the gene’s TSS (Pliner et al., 2018; Granja et al., 2021). Recent studies (Zhang et al., 2022; Bravo González-Blas et al., 2023) used machine learning methods, such as XGBoost (Chen et al., 2016), utilizing chromatin accessibility counts from multiple cRE candidates around the target gene as input to predict gene expression levels. These models identify cREs via contribution scores and, by incorporating multiple cREs, reduce false-positives and improve precision. However, models based solely on chromatin accessibility require separate training to predict the expression level of each target gene, increasing the risk of overfitting due to limited gene expression diversity relative to model parameters, and limits their ability to learn transcriptional regulation rules that apply across multiple genes.

Another machine learning approach focuses on the relationship between DNA sequences and the characteristics of cREs, such as quantitative TF binding (Avsec et al., 2021b), histone modification (Kelley et al., 2018; Avsec et al., 2021a; Yuan et al., 2022), and chromatin accessibility (Kelley et al., 2018; Avsec et al., 2021a; Yuan et al., 2022) or activity in reporter assays (de Almeida et al., 2022; de Almeida et al., 2024). While these models cannot be directly applied to single-cell-level cRE-gene relationships because they only accept DNA sequences as input, they suggest that the DNA sequence of cRE regions has sufficient information to predict cRE characteristics regardless of the interacting target gene. Thus, incorporating the DNA sequence of cRE candidates into the prediction of gene expression levels might enable the training of comprehensive models, that can learn gene-cRE relationships with a single parameter set and improve the prediction performance. However, such architecture is lacking.

Here, we propose an attention-based deep learning model that integrates chromatin accessibility, DNA sequence information, and genomic distance to learn a comprehensive genetic regulatory code. Recently, deep neural networks have offered a promising avenue for learning complex regulatory rules without prior knowledge. Among these, an attention method that allows flexible learning independent of the order or size of the input data has achieved remarkable results (Vaswani et al., 2023), particularly in natural language processing. We propose this attention-based deep neural network framework for detecting cRE activity at single-cell resolution.

## Results

### Overview of the framework

Our framework uses a single sample of single-cell Multiome ATAC-seq+GEX (scMultiome) data to simultaneously obtain single-cell RNA sequencing (scRNA-seq) and scATAC-seq information from the same cells. First, we trained deep neural networks to predict gene expression levels at the single-cell level from scATAC-seq counts, DNA sequences, and the genomic distance of the gene’s neighboring ATAC-seq peaks (Fig. 1A, left). Then, our framework calculated the contribution score of the cRE candidates to the expression of the target gene (Fig. 1A, right), which reflects the potential contribution of each peak to gene expression. Previous studies (Zhang et al., 2022; Bravo González-Blas et al., 2023) have reported that high contribution scores in regression models are associated with high cRE activity. Therefore, we used the contribution score as an indicator of cRE activity at the single-cell level.

**Fig. 1.**
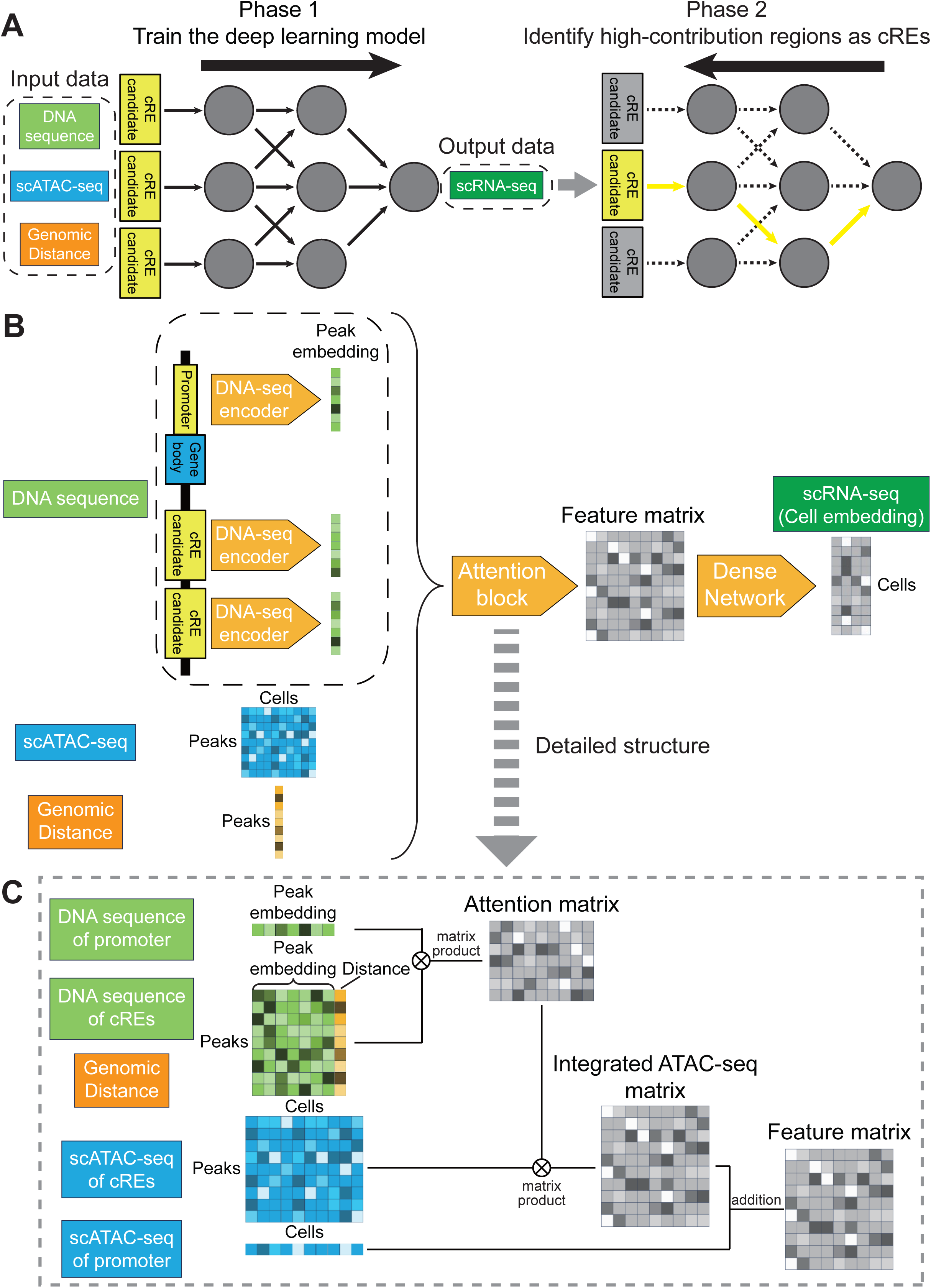
Workflow to detect cREs at the single-cell level. (A)The strategy to detect cRE regions using the deep learning method. As a first step, our framework trains a deep neural network to predict scRNA-seq counts from scATAC-seq counts, DNA sequence, and genomic distance. Next, the contribution of each cRE candidate to the gene expression level is calculated, and the regions with high contribution are determined as cREs. (B) The model structure of the attention-based neural network to predict gene expression level from DNA sequence, ATAC-seq, and genomic distance. First, the DNA-seq encoder extracts important features from the DNA sequence. Next, the features derived from the DNA sequence, scATAC-seq counts, and genomic distance are input into the Attention block. The Attention block integrates these input data and converts it into a feature matrix. Finally, the Dense network predicts gene expression level. (C) The detailed structure of the Attention block. First, the attention matrix is calculated between the promoter and cRE candidates based on the peak embedding derived from the DNA sequence and genomic distance. This attention matrix stores information on which cRE candidates have important features. By applying this attention matrix to scATAC-seq counts, an integrated scATAC-seq matrix is generated that only contains scATAC-seq counts for important regions. Finally, the scATAC-seq counts for promoters are added to generate a feature matrix. cRE, cis-regulatory elements; scRNA-seq, single-cell RNA sequencing; scATAC-seq, single-cell assay for transposase-accessible chromatin sequencing;

Our model assumes that gene expression levels are determined by the combination of gene promoter and cRE activities, based on previous knowledge indicating that the combination of promoter and enhancer activities can explain more than 60% of the variance in gene expression levels (Bergman et al., 2022). For each gene, we defined promoter peaks as ATAC peaks within ±500bp of the gene’s TSS and considered all ATAC peaks within ±300,000bp of the gene’s TSS as cRE candidates. The input data for the models comprised the DNA sequence, genomic distance, and scATAC-seq counts of the promoter and cRE candidates of each gene. The output was the normalized scRNA-seq count.

Our neural network models consist of the DNA-seq encoder, which extracts DNA sequence features from raw sequence data, and the Attention Block, which combines ATAC-seq, genomic distance, and the output of the DNA-seq Encoder (Fig. 1B). In our framework, the DNA-seq encoder first compressed all ATAC peaks one by one and extracted important motif information from the 1,344bp DNA sequence of all input peaks (Fig. 1B, upper left). This 8- layer convolutional neural network (CNN) compressed each peak to a 32-dimensional peak embedding.

The attention block had a structure similar to that of a general cross-attention-based neural network (Rombach et al., 2019) (Fig. 1C). Initially, it extracts features from peak embeddings and the genomic distance between the promoter and candidate cREs to compute the cross-attention between these elements (Fig. 1C, Attention Matrix). This attention matrix highlighted important cRE candidates based on DNA-seq data and genomic distances. In a typical cross-attention-based neural network, the next step is to apply the attention matrix to the same feature matrix used to calculate the cross-attention (in this case, the matrix of peak embeddings and genomic distance) to extract the relevant features. However, our model uniquely applied the attention matrix to the scATAC-seq count matrix, extracting the scATAC-seq features of important regions identified by DNA-seq features and genomic distance (Fig. 1C, integrated scATAC-seq matrix). This cross-modal attention mechanism enabled the integration of DNA sequences, genomic distances, and scATAC-seq information using a simple cross-attention structure. Finally, the scATAC-seq counts from the promoter regions were integrated to produce a final Feature Matrix (Fig. 1C).

Finally, the dense neural network converted the feature matrix from the attention block to the gene expression level (Fig. 1B, right). To stabilize the learning, we used principal component analysis (PCA) coordinates derived from scRNA-seq data instead of normalized gene expression. After learning the neural network model, our framework used DeepLIFT, which was developed as a tool to calculate the contribution score of a neural network (Shrikumar et al., 2019), to evaluate the contribution of cRE candidates and promoters to gene expression levels.

Model training consisted of two steps. First, the DNA-seq encoder was trained following a methodology similar to scBasset (Yuan et al., 2022). Specifically, a DNA-seq encoder was trained to predict chromatin accessibility at the single-cell level using DNA sequences. This process enabled the DNA-seq encoder to learn the critical sequence patterns for chromatin accessibility. Training of the DNA-seq encoder ran for 1,000 epochs. Second, the attention block and dense network were trained using scRNA-seq and scATAC-seq. 10% of the ATAC peaks and 20% of the cells were reserved as test data, and the remaining data were used for training.

### Benchmarking of cREs prediction performance

First, we benchmarked our framework using the public scMultiome dataset of human peripheral blood mononuclear cells (PBMC) from healthy donors (10x Genomics, 2021). The data includes 11,898 cells, 36,601 genes, and 143,887 peaks as raw data. After filtering, 11,754 cells, 6,853 genes, and 36,071 peaks were retained. The maximum number of peaks associated with a single gene was 79. After training the model and defining cRE activity as 60% cRE from the higher end, 83,519 cRE-gene pairs were detected. After annotating cell types based on their gene expression (Fig. 2A), our framework detected cell type-specific cRE activities, such as CD4^+^ T cell-specific (Fig. 2B, left) and CD8^+^ T cell-specific (Fig. 2B, right) cRE activities in the CD3D gene region, despite similar gene expression levels in these cell types (Fig. 2C).

**Fig. 2.**
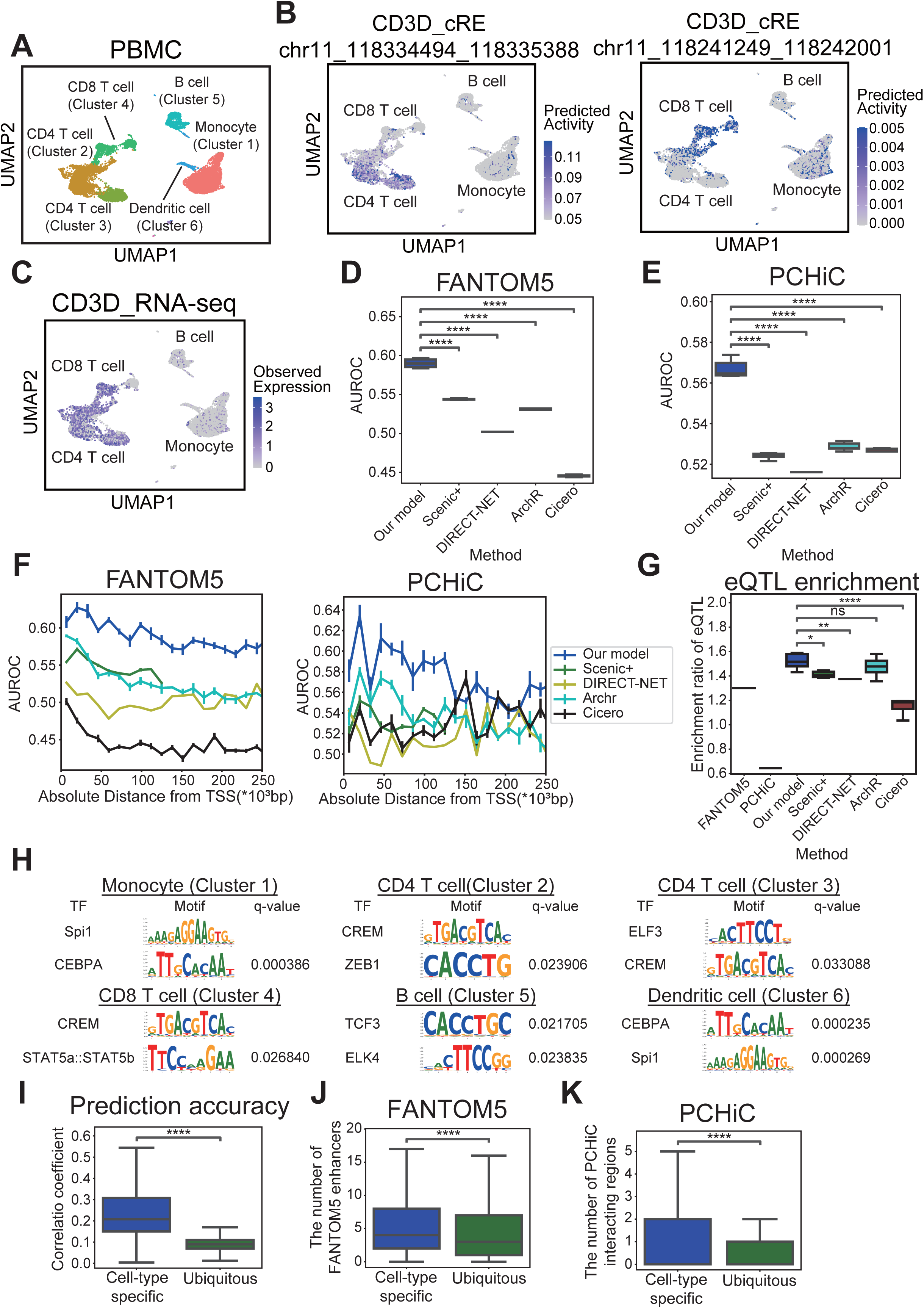
Model performance to predict cREs. (A)Cell type annotation of PBMC scMultiome data mapping was performed using scRNA-seq counts. (B, C) cREs (B) and observed gene expression levels of (C) CD3D. UMAP mapping was performed by scRNA-seq. Blue indicates gene expression level or predicted activity. (D, E) Prediction accuracy of cREs in the FANTOM5 (D) and PCHiC datasets (E). Each tool was trained 5 times. Boxplot represents quartile points. ns, 5.0*10^−2^ < p <= 1.0; *, 1.0*10^−2^ < p <= 5.0*10^−2^; **, 1.0*10^−3^ < p <= 1.0*10^−2^; ***, 1.0*10^−4^ < p <= 1.0*10^−3^; ****, p <= 1.0*10^−4^ (vs. our model, Student’s *t*-test, unpaired). (F) Prediction accuracy of cREs by distance from the TSS in the FANTOM5 dataset (left) and PCHiC data (right). Accuracy was measured using AUROC. The length of each bin was 1,500 bp. Error bars indicate standard deviations. (G) eQTL enrichment ratio of cREs predicted using computational tools, including our model and experimental methods. The box plot shows the result of 5-fold learning. Boxplot represents quartile points. ns, 5.0*10^−2^ < p <= 1.0; *, 1.0*10^−2^ < p <= 5.0*10^−2^; **, 1.0*10^−3^ < p <= 1.0*10^− 2^; ***, 1.0*10^−4^ < p <= 1.0*10^−3^; ****, p <= 1.0*10^−4^ (vs. our model, Student’s *t*-test, unpaired). (H) Enriched transcription factor-binding motifs in cell type-specific cREs. The enriched motifs and q-values were calculated using TF-MoDISCo (Shrikumar et al., 2020). (I) Comparison of the accuracy of the prediction of gene expression levels between genes with cell type-specific expression and those with ubiquitous expression. The y-axis shows Spearman’s correlation coefficient. p-value=0.000. Student’s t-test was used after Fisher’s Z-transformation to statistically test the correlation coefficients. (J, K) Comparison of the number of cRE regions linked to genes with cell type-specific and ubiquitous expression. (J) In the case of the FANTOM5 dataset, p=6.201*10^−18^ and (K) in the case of PCHiC data, p=1.770*10^−09^. Unpaired Student’s t-test was used for statistical analysis. cRE, cis-regulatory elements; PBMC, peripheral blood mononuclear cells; scMultiome, single-cell Multiome ATAC-seq+GEX; scRNA-seq, single-cell RNA sequencing; UMAP,Uniform Manifold Approximation and Projection; PCHiC, Promoter Capture Hi-C; AUROC, area under the receiver operating characteristic curve;

To assess how well our model performs at predicting cREs, we decided to benchmark it against existing tools. Cicero (Pliner et al., 2018) and ArchR (Granja et al., 2021) use scATAC-seq counts at candidate regions to detect cREs by correlating them either with scATAC-seq counts at promoter regions (Cicero) or with scRNA-seq gene counts (ArchR). On the other hand, DIRECT-NET (Zhang et al., 2022) and SCENIC+ (Bravo González-Blas et al., 2023) rely on XGBoost, a machine-learning method that predicts gene expression levels from scATAC-seq (Chen et al., 2016). To compare these different models, we then measured how well they performed at predicting cREs from the FANTOM5 database (Lizio et al., 2019) or detected using Promoter Capture Hi-C (PCHiC; Javierre et al., 2016). Importantly, our framework showed a significantly higher area under the receiver operating characteristic curve (AUROC) for both datasets (Fig. 2D, E), indicating higher accuracy.

The likelihood of cRE-gene interactions decrease with increasing genomic distances, and the risk of false positives is generally higher for distant cRE-gene pairs. While our model directly integrates distance information, Cicero and ArchR simply apply uniform penalties to distant cRE-gene pairs (Pliner et al., 2018; Granja et al., 2021), which might hinder the detection of bona fide long-range interactions. To assess how genomic distance affects our model’s predictions, we stratified gene-cRE pairs into 20 bins of increasing genomic distance. Importantly, our framework showed a higher prediction performance at every distance using CAGE-seq data (Fig. 2F, left) and at most distances using PCHiC data (Fig. 2F, right).

Next, we wanted to probe whether the cREs detected by our model were enriched for expression quantitative trait loci (eQTL), which would indicate that they correspond to bona fide cREs. We therefore used the eQTL database from the GTEx portal (GTEx Consortium, 2020) to show that our predicted set of cREs showed significantly higher eQTL enrichment ratios compared to cREs inferred using CAGE-seq or PCHiC contacts (Fig. 2G, p- value=1.89*10^−3^ for FANTOM5, p-value= 8.04*10^−6^ for PCHiC), suggesting that our model outperforms existing ones at detecting functional cRE-gene pairs.

### Functional characteristics of predicted cREs

Besides dectecting cRE-gene pairs, our framework evaluates DNA sequence motifs that are crucial for gene expression, allowing us to identify key TFs and infer transcriptional regulatory gene networks. After computing the contribution score of the input DNA sequence using Integrated Gradients (Sundararajan et al., 2017) with single-base pair resolution, we used TF-MoDISCo (Shrikumar et al., 2020) to identify the enriched DNA motif patterns contributing to gene expression levels. In the PBMC dataset, our framework successfully detected established cell type-specific TFs, such as CEBPA in monocytes (Scott et al., 1992), STAT5 in CD8^+^ T cells (Tripathi et al., 2010), and ELK4 in B cells (Yasuda et al., 2008), among the top five highly contributing DNA motifs (Fig. 2H).

In parallel, we interestingly observed that our model was generally more accurate at predicting the expression levels of cell type-specific genes compared to ubiquitously expressed housekeeping genes (Fig. 2I). A previous study (Zabidi et al., 2015) reported that cell type-specific genes are more dependent on distal cREs for their expression compared to housekeeping genes, which led us to hypothesize that differences in prediction accuracy might depend on the extent to which gene expression depends on cRE control. Consistently, we found that cell type-specific genes had a significantly larger number of cREs compared to housekeeping genes, using both the FANTOM5 or the PCHiC dataset (Fig. 2J, K).

### Analysis of tumor-specific cRE regulation

Next, we wanted to evaluate the performance of our model at predicting differences between cell types, by measuring their clustering performance using healthy human PBMC data (10x Genomics, 2021). The cells were clustered using k-means, either on the predicted cRE activity from our model or using eRegulon activity inferred using Scenic+ (Bravo González-Blas et al., 2023). Importantly, the cRE activities predicted by our model outperformed eRegulon predictions, as they showed a significantly higher Adjusted Rand Index (ARI, see EXPERIMENTAL PROCEDURES).

To further assess the capacitiy of each model at identifying functionally relevant cREs, we then used the top 15% cREs with the highest contributions scores detected by each model, and used overlapping scATAC-seq counts to cluster the different cell types present in the data. The cREs identified by our model clustered cells with a higher ARI compared to other tested tools (Fig. 3A, right panel), suggesting that it is more efficient at identifying relevant cREs from scATAC-seq data. However, ARI scores were consistently lower than those obtained using our model’s predicted cRE activity or the eRegulons from Scenic+ (Fig. 3A), indicating that scATAC-seq counts do not perfectly reflect the gene expression status of cells. However, our model can use this data to identify differences in cRE regulation between cell types that previous methods overlooked.

**Fig. 3.**
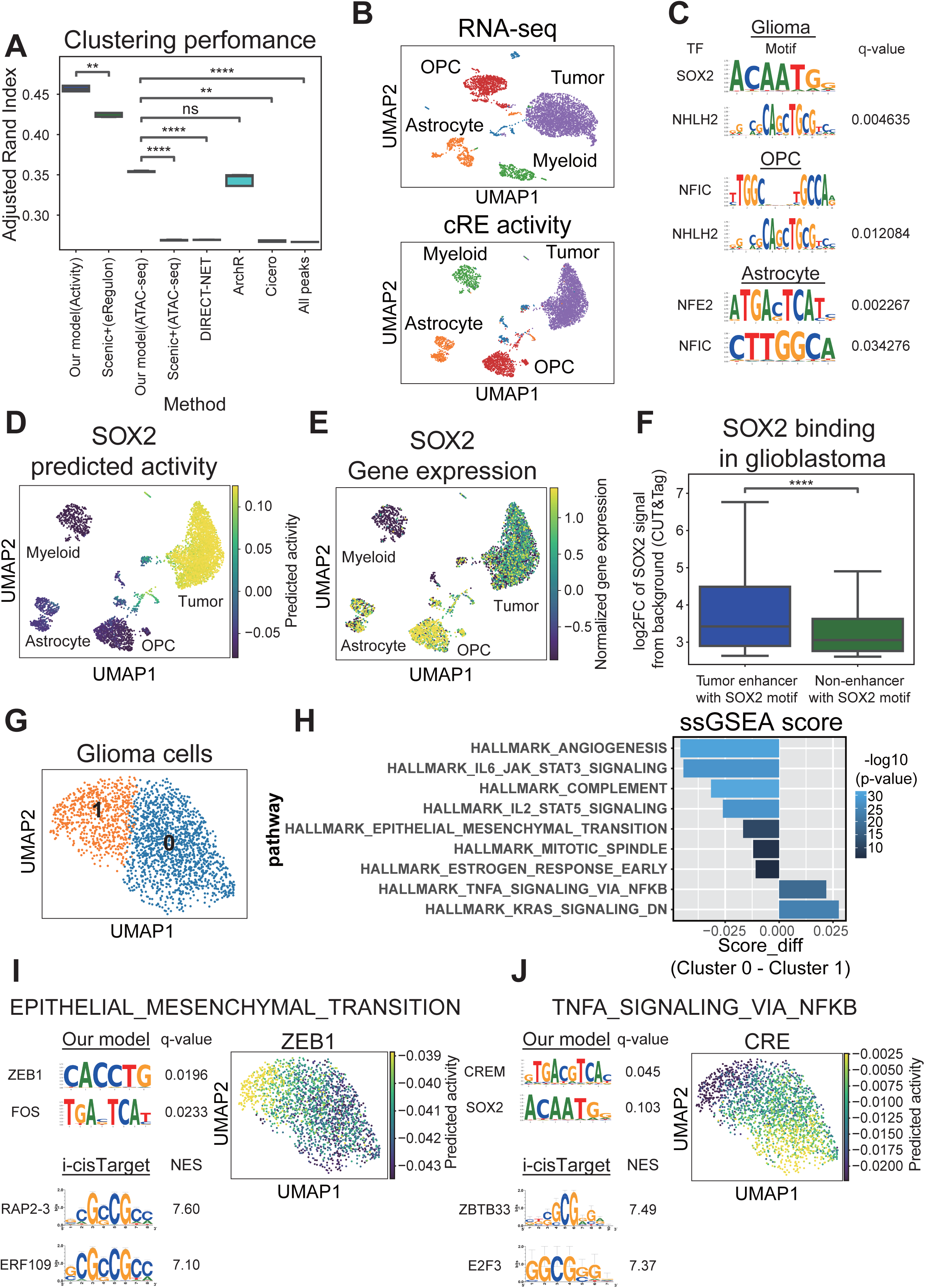
The analysis of tumor-specific cRE regulation. (A) Clustering performance in PBMC data. The Y-axis represents the adjusted Rand index. The first two plots on the left show the clustering performance of predicted cRE activity from our model and eRegulon activity from Scenic+ (Bravo González-Blas et al., 2023). The remaining plots compare clustering performance based on scATAC-seq counts, including the top 15% of high-activity peaks predicted by each tool and the total scATAC-seq counts. ns, 5.0*10^−2^ < p <= 1.0; *, 1.0*10^−2^ < p <= 5.0*10^−2^; **, 1.0*10^−3^ < p <= 1.0*10^−2^; ***, 1.0*10^−4^ < p <= 1.0*10^−3^; ****, p <= 1.0*10^−4^ (vs. our model, Student’s *t*-test, unpaired). (B) UMAP of glioma samples based on the scRNA-seq counts and predicted cRE activities. scRNA-seq-based UMAP was generated from the scRNA-seq counts of all genes. cRE activity-based UMAP was generated using the top 15% of cREs with the highest activity. Colors represent cell type annotations based on scRNA-seq counts. (C) The cell type-specific transcription factors detected by our model and TF-MoDISCo. Significant motifs, defined by a q-value of <0.05, were plotted for gliomas, OPC, and astrocytes. The top two motifs were highlighted in each case. (D) Predicted SOX2 activity at the single-cell level. The color indicates the predicted SOX2 activity. (E) SOX2 gene expression levels in scRNA-seq counts. Colors indicate log-normalized and scaled gene expression levels. (F) SOX2 CUT&Tag signaling between tumor-specific cREs and non-cRE regions with a SOX2 binding motif. The Y-axis represents the log2 fold change in the SOX2 CUT&Tag signal from glioma cell samples relative to the background signal. p=2.4*10^−10^ (Student’s *t*-test, unpaired). (G) Clustering of glioma cells based on predicted cRE activity. The UMAP is based on the cRE activities. (H) Enrichment of gene sets in intra-tumor clusters by ssGSEA analysis. Statistical analysis was performed using the Mann–Whitney U test. (I, J) Gene sets and transcription factors involved in the significant enrichment of each intra-tumor cluster. The motif enrichment analysis was performed using TF-MoDISCo (Shrikumar et al., 2020) and i-cisTarget (Herrmann et al., 2012). (I) Results for “HALLMARK_EPITHELIAL_MESENCHYMAL_TRANSITION” and (J) “HALLMARK_TNFA_SIGNALING_VIA_NFKB.” cRE, cis-regulatory elements; PBMC, peripheral blood mononuclear cells; scRNA-seq, single-cell RNA sequencing; scATAC-seq, single-cell assay for transposase-accessible chromatin sequencing; UMAP, Uniform Manifold Approximation and Projection; OPC, oligodendrocyte precursor cells;

Next, to verify whether our model can detect tumor-specific cRE regulation, we applied our model to public single-cell data from patients with pediatric glioma (Jessa et al., 2022; 66,070 cREs detected in total, see EXPERIMENTAL PROCEDURES). First, cell clusters identified using scRNA-seq and cRE activity corresponded to the cell types identified by scRNA-seq (Fig. 3B). To determine the key DNA binding motifs associated with cell type-specific gene expression changes, we then performed motif enrichment at cell type-specific cREs using TF-MoDISCo (Shrikumar et al., 2020). This way, we identified sequence signatures for the NHLH2 and NFIC neuronal factors (Frazel et al., 2023; Wilczynska et al., 2009) but also for SOX2 (Fig. 3C), which has been associated with poor prognosis in gliomas (Garros-Regulez et al., 2016) and was the most enriched motif in glioma-specific cREs. Consistent with motif analysis, predicted SOX2 activity in cREs was significantly higher in glioma cells (Fig. 3D, p=0.000). However, SOX2 was more highly expressed in oligodendrocyte precursor cells (OPCs) than in glioma cells (Fig. 3E), highlighting the limitation of relying solely on TF gene expression levels to predict TF activity.

To further validate SOX2 binding to tumor-specific cREs, we used publicly available SOX2 CUT&Tag data from patients with glioma (Benedetti et al., 2022). As predicted by our model, tumor-specific cREs with SOX2 binding motifs showed significantly higher CUT&Tag signals than non-cRE regions with the SOX2 motif (Fig. 3F, p=2.4*10^−10^). This result demonstrates that our model can effectively integrate DNA sequence, epigenetic, and transcriptomic information to predict TF-binding events, which is difficult to achieve using only DNA sequence data.

Finally, we analyzed the intra-tumor heterogeneity of cRE regulation. After clustering tumor cells into two groups using k-means on predicted cRE activity (Fig. 3G), we examined their differences in gene expression levels using ssGSEA (Subramanian et al., 2005). Cluster 1 showed elevated expression of angiogenesis- and epithelial-mesenchymal transition (EMT)- related genes (Fig. 3H). Focusing on EMT-related genes, we identify TFs regulating the cREs associated with these genes by our model with TF-MoDISCo (Shrikumar et al., 2020) (Fig. 3I, left). ZEB1, a well-known EMT-related TF (Zhang et al., 2015), emerged as a promising candidate. Consistent with this, we observed a gradual increase in ZEB1 predicted activity from Cluster 0 to Cluster 1 (Fig. 3I, right), while existing statistical methods (i-cisTarget; Herrmann et al., 2012) failed to identify significant motifs in the EMT-related cREs (Fig. 3I, bottom-left). Thus, our model can detect intra-tumor heterogeneity in EMT-related cell states and identify key TFs that regulate EMT-related genes.

On the other hand, genes involved in TNF-α and NFκB signaling pathways were more expressed in cluster 0. However, when analyzing the cREs associated with these genes using TF-MoDISCo, the binding motif for NFκB was not enriched. Instead, CREM, a TF belonging to the cAMP response element (CRE) family, was identified as the top motif (Fig. 3J, left). CREB1, a member of the CRE family, has been reported to co-regulate NFκB target genes with RelA (Nakayama et al., 2013). Consistent with this finding, the predicted CRE activity was specific to cluster 0 (Fig. 3J, right). In conclusion, our model was able to pinpoint subpopulation-specific TF signatures within tumors, using single-cell multi-omics data.

## Discussion

This study proposed a new method for detecting cRE activity at the single-cell level using deep neural networks equipped with attention mechanisms. Our model integrates DNA sequence information, chromatin accessibility, and genomic distance within the model architecture, leading to a more accurate detection of cRE activity compared to previous methods. Notably, our model learns transcriptional regulation rules in a data-driven manner by incorporating genomic distance from the TSS, which is manually included in other models (Pliner et al., 2018; Granja et al., 2021). This data-driven approach allows the model to handle distance information more effectively than rule-based methods, such as Cicero and ArchR (Pliner et al., 2018; Granja et al., 2021).

Additionally, our model could detects differences in cRE regulation between cell types more accurately than previous tools. In our analysis of glioma samples, the model accurately identified the SOX2 transcriptional activity specific to gliomas. Additionally, intra-tumor analysis revealed heterogeneity in the regulation of EMT-related genes, which are key factors in metastasis. The model also identified the critical TFs involved in the regulation of these cREs. When tumors acquire metastatic abilities or become resistant to treatment, only a subset of tumor cells develops these characteristics (Dagogo-Jack et al., 2018). Additionally, in certain cancers, such as acute myeloid leukemia, only a fraction of cells, such as leukemia stem cells, possess the capacity to regenerate large numbers of tumor cells (Shlush et al., 2014). Therefore, to assess tumor pathology and identify precise therapeutic targets, it is crucial to measure transcriptional regulation in specific cell subpopulations. Our model enables high-resolution analysis of specific gene sets within defined cell populations, making it a valuable tool for multi-omics single-cell data analysis.

In addition, our model was designed to analyze a smaller size of clinical samples of any disease in a hospital setting without requiring pre-training on large datasets. This allowed us to assess cRE activity at the single-cell level, providing insights into the heterogeneity of cRE regulation in rare diseases. However, a limitation of our method is that deep learning-based regression models capture only the correlations between inputs and outputs. Thus, a high contribution score to the gene expression level does not guarantee a causal relationship between the detected cREs and target genes. To address this, one potential solution involves training the model on a large dataset and using causal analysis approaches such as a causal transformer (Melnychuk et al., 2022). Another approach involves experimental validation. Nevertheless, our model prioritizes versatility and applicability to single samples by relying on extensive pre-training on large datasets. Another limitation is that it may be difficult to detect cRE activity that is consistently high in all cells. As this model relies on differences in activity and gene expression levels between cells, it is expected that it will not be possible to learn about cREs that do not differ between cells. To address this issue, analysts should carefully select appropriate reference cell types based on the analysis goal and include these cells in the samples.

## EXPERIMENTAL PROCEDURES

### Preparation of scMultiome

The PBMC dataset comprised “Peripheral Blood Mononuclear Cells from a Healthy Donor, with Granulocytes Removed Through Cell Sorting (10k),” sourced from the 10x website (10x Genomics, 2021). We analyzed the respective available count data, namely “pbmc_granulocyte_sorted_10k_filtered_feature_bc_matrix.h5” for PBMC datasets. Subsequently, scRNA-seq and scATAC-seq data were processed using Scanpy (version 1.9.3) (Wolf et al., 2018). Only genes expressed in 5% or more of the cells and cells expressing 500 or more genes were included for downstream analysis.

### Annotation of cells for PBMCs

Cell clustering was conducted using Seurat (version 4.3.0) (Hao et al. 2021) with the parameter “resolution=0.1.” The cells were annotated using marker genes. We used S100A8 and CD14 for monocytes, CD3D and CD4 for CD4^+^ T cells, PRF1 for CD8^+^ T cells, CD19 and EBF1 for B cells, and CD1C for dendritic cells.

### Prediction model for gene expression levels

Our framework was designed to use single sample scMultiome data as the input. For each gene, the TSS position in the GRCh38.p13 genome was obtained from the Ensembl database using BiomaRt (version 2.50.3) (Durinck et al., 2005). First, ATAC-seq peaks located within ±300,000 bp of the TSS were considered candidate cRE regions for the respective gene. The peak located within ±500bp of the TSS of the genes was identified as the promoter peak. In cases where multiple candidate promoter peaks fell within this range, the closest peak was defined as the promoter peak. The promoter peak of each gene was excluded from the candidate cRE region. Genes lacking promoter peaks or cRE candidate regions were excluded from the downstream analysis. The model was trained on a per-gene basis, with each gene constituting a single dataset. Specifically, the input consisted of chromatin accessibility counts for the cRE candidate region/promoter, DNA sequence, and distance from the TSS, whereas the output was normalized to gene expression. Our gene expression prediction model comprised two components: a DNA sequence encoder and an attention-based block. The DNA sequence encoder was an 8-layer CNN that accepted a one-hot vector representation of a 1,344 bp nucleotide sequence as input and yielded a compressed 32- dimensional representation. The attention-based block initially processed the 32-dimensional DNA sequence features of cRE candidate regions and promoters and 1-dimensional distance information through a layer of a Feed Forward Neural Network (FFNN), followed by the computation of an attention matrix via the matrix product. Subsequently, the product of the attention matrix and ATAC-seq counts was used to integrate the ATAC-seq information with the DNA sequence information. The resulting matrix was normalized using Instance Normalization. The ATAC-seq information of the promoter was subsequently added and a feature matrix was computed. Finally, the gene expression levels were derived from the feature matrix using a two-layer FFNN.

### Model training

First, the dataset was divided into training and testing datasets. Initially, 10% of the input ATAC-seq peaks were earmarked as test data, whereas the remaining 90% constituted the training set. Only genes with all the cREs and promoters included in the training peaks were designated as training genes; the remainder were allocated to the test set. In terms of cells, 80% were designated as training cells and the remaining 20% were assigned to the test set. The initial step involved training the DNA sequence encoder using a method similar to that used in the scBasset (Yuan et al., 2022). Specifically, we extracted a 32-dimentional peak embedding from the DNA sequence and trained the model to predict ATAC-seq counts at the single-cell level based on these features. Pre-training was conducted for 1,000 epochs. The outputs from the DNA-seq encoder, such as 32-dimentional feature vectors and predicted ATAC-seq counts, were used for analyzing the peak embedding and noise-suppressed ATAC-seq counts, respectively. Subsequently, the scRNA-seq count matrix was compressed using PCA with the PCA transition matrix derived solely from the training cells. This compression reduced the scRNA-seq count matrix to 50 dimensions. Attention-based block training was then performed using a 32-dimensional compressed sequence representation, ATAC-seq, and distance information as input. The model was trained to predict 50-dimensional cellular coordinates compressed using PCA at the single-cell level. The model training was run for 500 epochs, with early stopping if the test loss did not improve after 10 consecutive epochs. After training, the predicted gene expression levels were calculated by inverting the PCA coordinates. Notably, to evaluate the prediction accuracy, a 10-fold cross-validation was conducted.

### Acquisition of cRE activity

cRE activity was determined by assessing the contribution of the input ATAC-seq count to gene expression levels. Contribution scores were calculated using the DeepLift function in the Captum package (version 0.6.0) (Kokhlikyan et al., 2020). For analyses other than benchmarking of cRE region prediction, we only used regions with a positive correlation between ATAC-seq counts and cRE activity to unify the interpretability of the results.

### Detection of cell type-specific cREs

The cell-type specificity of each cRE was determined by comparing its activity in a given cell type to its activity across all other cell types. Statistical significance was assessed using the Mann–Whitney U test. Peaks that met the criteria of adjusted p-value < 0.0001 and a mean activity difference greater than 0.01 were classified as cell-type-specific cREs.

### Motif analysis using the proposed model

To detect cell type-specific TF-binding motifs, we first identified cell type-specific genes. This was performed using Seurat (version 4.3.0) (Hao et al. 2021), where differentially expressed genes (DEGs) were defined as those with a log2 fold change greater than 0 and p-value < 0.05 when comparing the target clusters to other clusters. Cell type-specific cREs related to these DEGs were used for motif analysis. In order to suppress the noise in the contribution score, only cells in which the cREs linked to that gene are constantly active were used in the analysis, and cells in which cRE activity was predicted to be high by chance were excluded. Then, the cells having top 25% cRE activities in over 70% of cREs related to the target gene are included in the analysis. This analysis was performed using TF-MoDISCo (lite version 1.0.0) (Shrikumar et al., 2020), which utilized the contribution scores calculated using our model. The position weight matrices (PWMs) identified by TF-MoDISCo were then compared with the JASPAR CORE 2024 vertebrate (non-redundant) database (Rauluseviciute et al., 2024). TF motifs with a q-value less than 0.05 were considered statistically significant. When searching for motifs related to a certain gene set, we used the target gene set instead of DEGs.

### Evaluation of cRE prediction results

The cRE prediction results were evaluated using data from the FANTOM5 database (Lizio et al., 2019) and PCHiC data (Javierre et al., 2016) for PBMCs. The enhancer list of the FANTOM5 database was derived from “F5.hg38.enhancers.bed.gz”, which is the curated list of enhancers in the FANTOM5 database. PCHiC data were obtained from “PCHiC_peak_matrix_cutoff5.tsv.” Among the list of ATAC-seq peaks used as model inputs, regions that overlapped by more than a single base pair with FANTOM5 enhancers were considered FANTOM5 enhancer regions. As with the analysis of the FANTOM5 dataset, we calculated the overlap between the ATAC-seq peaks used as input for the model and PCHiC peaks. The cRE candidates in which the promoter region of PCHiC overlapped with the model promoter region and the other end of PCHiC overlapped with the cRE candidate region were identified as cRE regions. The predictive performance of our model was compared with that of Cicero, ArchR, DIRECT-NET, and Scenic+ (Pliner et al., 2018; Granja et al., 2021; Zhang et al., 2022; Bravo González-Blas et al., 2023). Each detected cRE-gene interaction was identified according to the respective tutorial of the tool. To compare performance, we calculated the overlap of cRE-gene pairs detected by each tool with the ATAC-seq peak regions that served as the input to our model. The overlapping ATAC-seq peak gene pairs were considered as the cRE-gene pairs detected by each tool. Prediction performance was quantified as the AUROC using the roc_auc_score function in sklearn (version 1.4.0) (Pedregosa et al., 2011). To calculate the AUROC per distance, the distance from the TSS was divided into 20 bins ranging from 0 to 28,000 bp. AUROC was then calculated for the cRE-gene pairs within each bin.

### Processing of eQTL data

The eQTL data were obtained from the GTEx v8 database (GTEx Consortium, 2020). Only variants with PIP > 0.5 were defined as causal variants for those detected in whole blood cells. The enrichment ratio was calculated as follows (Sakaue et al., 2024):

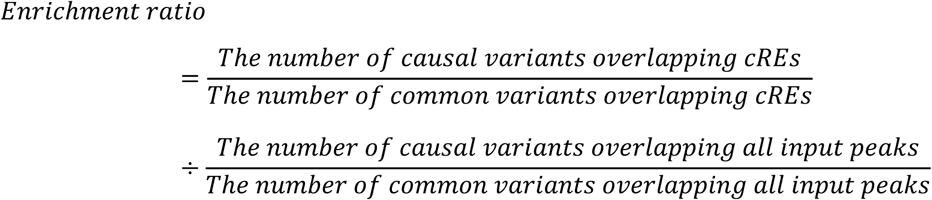

### Calculating the distance from the TSS

The TSS positions of all genes in the GRCh38.p13 genome were obtained from the Ensembl database using the biomaRt package (version 2.50.3) (Durinck et al., 2005). Distances between all candidate cREs and the TSS of the closest gene were calculated.

### Analysis of cell type-specific genes and genes with ubiquitous expression

Genes with cell type-specific expression were selected using Seurat (version 4.3.0) (Hao et al. 2021), with the criteria of log2 FC>0 and p-value <0.05 in each cluster. Genes not included in the set of genes with cell-type-specific expression were defined as those with ubiquitous expression. The accuracy of the gene expression prediction was evaluated using Spearman’s correlation coefficient. Fisher’s Z-transformation was used for the statistical test of the correlation coefficient, and Student’s t-test was used.

### Benchmarking of clustering performance

Clustering performance was benchmarked using ARI, with scRNA-seq-based clustering serving as the ground truth. The standard Seurat pipeline (version 4.3.0) (Hao et al. 2021) was used for scRNA-seq clustering. Specifically, following normalization, the top 2,000 variable features were selected for PCA. Clustering was then performed using PCA coordinates with the resolution parameter set to 0.1, resulting in the identification of nine distinct clusters. For clustering based on the predicted cRE activity from our model, the top 15% of the regions with the highest predicted cRE activities were selected, and k-means clustering was performed to group the cells into nine clusters based on these predicted activities. In the case of eRegulon clustering from Scenic+ (Bravo González-Blas et al., 2023), the “eRegulon_AUC” values generated by Scenic+ were used for clustering. For clustering based on scATAC-seq counts, the top 15% of regions showing the highest activity, as identified by each computational tool, were used. Cells were grouped into nine clusters using k-means clustering. For the clustering based on “all peaks,” all scATAC-seq peaks were included in the analysis. ARI was calculated using the “adjusted_rand_score” function, and k-means clustering was conducted using the “KMeans” function from the sklearn package (version 1.4.0) (Pedregosa et al., 2011). Statistical comparisons were made using the Student’s t-test, and differences with p-value < 0.05 were considered statistically significant.

### Analysis of the sample from patients with glioma

We obtained the scMultiome data for a glioma sample from NCBI GSE210568 (Jessa et al., 2022). Specifically, we analyzed the sample labeled “P-1694_S-1694_multiome.” Data were processed using the same methodology that was applied to PBMCs, including the prediction of cRE activity. The raw data included 5,530 cells, 60,658 genes, and 107,873 peaks. After filtering, the dataset for analysis comprised 5,304 cells, 7,805 genes, and 33,266 peaks. The maximum number of peaks associated with a single gene was 69. Uniform Manifold Approximation and Projection (UMAP) visualization was generated based on the predicted activity of the top 15% of the most active cREs using Scanpy (version 1.9.3) (Wolf et al., 2018).

### Cell annotation for the glioma sample

Cell annotation was performed based on the marker genes reported in the existing literature. Specifically, PDGFA and FGFR2 for gliomas (Verhaak et al., 2010; Jimenez-Pascual et al., 2019), PDGFRA for OPCs (Pringle et al., 1992), GFAP for astrocytes (Hol et al., 2015), and S100A8 for myeloid cells (Odink et al., 1987) were used.

### Detection of transcription factor binding regions

The cREs bound by each TF were identified using the FIMO function from the MEME Suite (version 5.5.5) (Bailey et al., 2015), with PWM files sourced from the JASPAR CORE 2024 vertebrate (non-redundant) database. FIMO was run with a threshold of 0.001 (’--threshold 0.001’). The top 5,000 highest-scoring matches were selected and defined as TF-binding regions.

### Calculation of transcription factor activity at the single-cell level

The predicted activity of TFs at the single-cell level was calculated as the average activity of cREs containing TF-binding motifs in each cell.

### SOX2 CUT&Tag analysis

The SOX2 CUT&Tag data were obtained from NCBI GSE200062 (Benedetti et al., 2022). Specifically, the file “GSM6008250_SMNB19_SOX2_3_broadPeak.bed.gz” was used for the analysis. From the results of our model, we identified tumor-specific cREs and non-cRE regions containing SOX2 binding sites for subsequent analysis. Overlapping SOX2 CUT&Tag broad peaks were detected for each group. To detect these overlapping peaks, we used the intersect function of Bedtools (version 2.30.0) (Quinlan et al.,2010) with the parameters “-wa -u -F 1 -a cRE regions from models -b SOX2 CUT&Tag peaks files”. The average log2 fold change from the background signal was calculated for each SOX2 CUT&Tag peak group. Statistical significance was evaluated using Student’s t-test, with p-value < 0.05.

### Clustering of tumor cells

The top 15% of cREs with high predicted activity were selected to map glioma cells using UMAP. The glioma cells were then clustered using the sc.tl.leiden function in Scanpy (version 1.9.3) (Wolf et al., 2018), with a resolution parameter of 0.2, resulting in two distinct clusters.

### ssGSEA analysis

First, to identify the gene groups regulated by cREs, we performed gene ontology analysis on the target genes of the top 15% of the most active predicted cREs using the H collection from the Molecular Signatures Database (MSigDB) (Liberzon et al., 2015). The “msigdbr” package (version 7.5.1) (Dolgalev et al., 2022) in R was used to obtain the gene sets. Gene sets with an adjusted p-value < 0.05 were considered to be regulated by cREs. Next, we used the ssGSEA package to assess the enrichment of these cRE-regulated gene sets in glioma cells and calculated the enrichment score at the single-cell level. We then used the Mann–Whitney U test to identify cluster-specific gene sets, with those having a p-value < 0.05 deemed significant. Finally, we calculated the differences in the average enrichment scores between the clusters.

### Transcription factor analysis of gene sets

First, cREs associated with the gene sets “HALLMARK_EPITHELIAL_MESENCHYMAL_TRANSITION” and “HALLMARK_TNFA_SIGNALING_VIA_NFKB” were identified. The contribution scores for these cREs were calculated at the single-nucleotide level, followed by motif analysis using TF-MoDISCo (lite version 1.0.0) (Shrikumar et al., 2020). TF motifs were then identified by comparing them with the JASPAR CORE 2024 vertebrate (non-redundant) motif database using a significance threshold of q-value < 0.05.

To complement this, conventional motif enrichment analysis was performed on the cRE regions corresponding to each gene set using i-cisTarget (Herrmann et al., 2012). Finally, the predicted TF activities of ZEB1 and CREM in the glioma cells were calculated and visualized as cRE contribution scores.

## ACKNOWLEDGMENTS

We would like to thank Dr. Hidetoshi Shimodaira (Kyoto Univ.), Dr. Makoto Taiji (RIKEN), Dr. Masako Iwasaki (Osaka Metropolitan Univ. & Osaka Univ.), Dr. Hajime Nagahara (Osaka Univ.), Dr. Yuta Nakashima (Osaka Univ.), Dr. Hideaki Hayashi (Osaka Univ.), Dr. Naoki Hosen (Osaka Univ.), Dr. Michiko Ichii (Osaka Univ.) and all members of JST CREST Bio-DX for their meaningful suggestions and discussion. We would like to thank Dr. Vincent Loubiere (Research Institute of Molecular Pathology) for helpful discussions and advice regarding the manuscript.

## AUTHOR CONTRIBUTIONS

K.M., K.I., and M.O. conceived the project. K.M. and K.I. designed the model. K.M. performed all the analyses. K.M., K.I., and M.O. interpreted the data and wrote the manuscript. M.O supervised the study. All the authors have read and approved the final version of the manuscript.

## ETHICS STATEMENT

Not applicable.

## DATA AVAILABILITY

The PBMC scMultiome dataset, “Peripheral Blood Mononuclear Cells from a Healthy Donor, with Granulocytes Removed Through Cell Sorting (10k)” was obtained from the 10x website (https://www.10xgenomics.com/datasets/pbmc-from-a-healthy-donor-granulocytes-removed-through-cell-sorting-10-k-1-standard-2-0-0). The enhancer list from the FANTOM5 database, “F5.hg38.enhancers.bed.gz,” was obtained from FANTOM5 website (https://fantom.gsc.riken.jp/5/). The promoter capture HiC (PCHiC) data, “PCHiC_peak_matrix_cutoff5.tsv,” was obtained from the Supplemental Information of Javierre et al. (2016) (“Data S1”; https://www.cell.com/cms/10.1016/j.cell.2016.09.037/attachment/5bc79f6f-1b69-4192-8cb8-4247cc2e0f39/mmc4.zip). The scMultiome data for a glioma sample “P-1694_S- 1694_multiome” was obtained from NCBI GSE210568 (https://www.ncbi.nlm.nih.gov/geo/query/acc.cgi?acc=GSE210568). SOX2 CUT&Tag data, “GSM6008250_SMNB19_SOX2_3_broadPeak.bed.gz” was obtained from NCBI GSE200062 (https://www.ncbi.nlm.nih.gov/geo/query/acc.cgi?acc=GSE200062). The code used in this study is available at https://github.com/okadalabipr/cREscENDO

